# MECP2 regulates cortical plasticity underlying a learned behavior in adult female mice

**DOI:** 10.1101/041707

**Authors:** Keerthi Krishnan, Billy Y. B. Lau, Gabrielle Ewall, Z. Josh Huang, Stephen D. Shea

**Author notes:** Corresponding author: Keerthi Krishnan. Author Contributions: KK and BYBL contributed equally. SDS and ZJH supervised the project. KK, BYBL, and SDS designed the experiments and developed the methods. KK, BYBL, GE and SDS collected and analyzed the data. KK, BYBL, ZJH and SDS wrote and edited the paper.

## Abstract

Neurodevelopmental disorders begin with the emergence of inappropriate synaptic connectivity early in life, yet how the sustained disruption of experience-dependent plasticity aggravates symptoms in adulthood is unclear. Here we used pup retrieval learning to assay adult cortical plasticity in a female mouse model of Rett syndrome (*MeCP2^het^*). We show that auditory cortical plasticity and retrieval learning are impaired in *MeCP2^het^*. Specifically, normal MECP2 expression in the adult auditory cortex is required for efficient retrieval learning. In wild-type mice, cohabitation with a mother and her pups triggered transient changes to auditory cortical inhibitory networks, including elevated levels of the GABA-synthesizing enzyme GAD67. However, *MeCP2^het^* further exhibited increased expression of parvalbumin (PV) and perineuronal nets (PNNs), events thought to suppress plasticity at the closure of critical periods and in adult learning. Averting these events with genetic and pharmacological manipulations of the GABAergic network restored retrieval behavior. We propose that adult retrieval learning triggers a transient episode of inhibitory plasticity in the auditory cortex that is dysregulated in *MeCP2^het^*. This window of heightened sensitivity to social sensory cues reveals a role of *MeCP2* mutations in facilitating adult plasticity that is distinct from their effects on early development.

## INTRODUCTION

Rett syndrome (RTT) is a genetic neuropsychiatric disease predominantly caused by mutations in the X-linked gene methyl CpG-binding protein 2 (*MeCP2*) [1]. Males with mutations of their lone copy of the gene suffer neonatal encephalopathy and die in infancy [2]. Most surviving RTT patients are females that are heterozygous for *MeCP2* mutations (*MeCP2^het^*). In these females, random X-chromosome inactivation leads to mosaic wild type MECP2 expression and consequently a syndromic phenotype. Patients achieve expected early postnatal developmental milestones, but later experience an abrupt regression around 6 - 12 months [3, 4]. They typically survive into middle age [5], exhibiting sustained sensory, cognitive and motor deficits throughout life.

MECP2 is broadly expressed in the developing and adult brain [6, 7] and is continually required to maintain adult neural function [8-10]. Moreover, restoration of normal MECP2 expression in adult mice improves symptoms [8-10]. These observations establish that MECP2 is necessary and sufficient to regulate brain function in adulthood. However, the specific ongoing function of MECP2 in the mature brain, apart from its widely studied role in early development, remains unclear.

MECP2 regulates neuronal chromatin architecture and gene transcription [11-13] in response to neural activity and experience during postnatal life [14, 15]. The known cellular function of MECP2 and the characteristic timing of disease progression raise the possibility that the regulation of neural circuits by MECP2 is magnified during specific windows of enhanced sensory and social experience throughout life. We therefore hypothesized that continued disruptions of experience-dependent plasticity in *MeCP2* mutants hinder adult learning. We tested this hypothesis in adult female mice using a natural, learned maternal behavior, the pup retrieval behavior, which is known to involve experience-dependent auditory cortical plasticity [16-18]. First-time mother mice respond to their pups’ ultrasonic distress vocalizations (USVs) by gathering the pups back to the nest, a critical aspect of maternal care [19, 20]. Virgin females with no previous maternal experience (‘surrogates’ or ‘Sur’) can acquire this behavior when co-housed with the mother and her pups [16]. Both in these ‘surrogates’ and in mothers, proficient pup gathering behavior is correlated with physiological plasticity in the auditory cortex [16-18]. Thus, the auditory cortex during maternal experience appears to be a location and time of heightened adult plasticity.

We report that adult *MeCP2^het^* surrogates and surrogates lacking MECP2 only in the adult auditory cortex exhibit impaired pup retrieval behavior. Maternal experience triggered changes in GABAergic interneurons of wild-type surrogates, but additional changes were observed in *MeCP2^het^* surrogates. Specifically, we observed elevated expression of parvalbumin (PV) and perineuronal nets (PNNs). Increases in expression of these markers are associated with the termination or suppression of plasticity in development and adulthood [21-26]. Genetic manipulation of GAD67, the primary synthetic enzyme for GABA, suppressed increases in PV and PNNs and restored gathering in *MeCP2^het^*. Furthermore, specific removal of the PNNs through chondroitinase ABC injections into the auditory cortex also restores efficient pup retrieval behavior in *MeCP2^het^*. Together, our results show that MECP2 regulates adult experience-dependent plasticity required for efficient gathering behavior through changes to the GABAergic interneuron network. They further suggest that dysregulated MECP2 expression leads to constriction of plasticity in response to experience, and that when this brake is removed, learning improves.

## MATERIALS AND METHODS

### Animals

All experiments were performed in adult female mice (7-10 weeks old) that were maintained on a 12-h–12-h light-dark cycle (lights on 07:00 h) and received food *ad libitum*. Genotypes used were CBA/CaJ, *MeCP2^het^* (C57BL/6 background; B6.129P2(C)-*Mecp2^tm1.1Bird^*/J), *MeCP2^wt^* and *MeCP2^flox/flox^* (B6.129S4-*Mecp2^tm1Jae^*/Mmucd). *MeCP2^flox/flox^* mice were bred with H2B-GFP (*Rosa26-loxpSTOPloxp-H2BGFP*) line [27] to facilitate identification of injected cells. The double mutant *MeCP2^het^; Gad1^het^* (*Het;Gad1^het^*) was generated by crossing *MeCP2^het^* females and *Gad1^het^* males. The *Gad1^het^* allele was generated using homologous recombination in ES cells; a cassette containing de-stabilized GFP cDNA (D2GFP) was inserted at the translation initiation codon (ATG) of the *Gad1* gene. The goal was to generate a *Gad1* gene transcription reporter allele, but the same allele is also a gene knockout. This design was essentially the same as the widely used *Gad1-GFP* knockin allele [28]. Targeted ES clones were identified by PCR and southern blotting. Positive ES clones were injected into C57BL/6 mice to obtain chimeric mice following standard procedures. Chimeric mice were bred with C57BL/6 mice to obtain germline transmission. D2GFP expression was weak and was restricted to GABAergic neurons throughout the mouse brain, indicating successful gene targeting. The colony is maintained as heterozygotes, as homozygotes are lethal. All procedures were conducted in accordance with the National Institutes of Health’s *Guide for the Care and Use of Laboratory Animals* and approved by the Cold Spring Harbor Laboratory Institutional Animal Care and Use Committee.

### Pup retrieval behavior and movement analysis

We housed two virgin female mice (one control and one experimental mouse; termed ‘surrogates’) with a first-time pregnant CBA/CaJ female beginning 1-5 days before birth. Pup retrieval behavior was assessed starting on the day the pups were born (postnatal day 0; D0) as follows: 1) one female was habituated with 3-5 pups in the nest of the home cage for 5 minutes, 2) pups were removed from the cage for 2 minutes and 3) one pup was placed at each corner and one in the center of the home cage (the nest was left empty if there were fewer than 5 pups). Each adult female had maximum of 5 minutes to gather the pups to the original nest. After testing, all animals and pups were returned to the home cage. The same procedure was performed again at D3 and D5. All behaviors were performed in the dark, during the light cycle (between 10:30 AM and 4:00 PM) and were video recorded. For analysis, a blind experimenter counted the number of errors and measured the latency of each mouse to gather all five pups. An error was scored for each instance of gathering of pups to the wrong location or of interacting with the pups (e.g. sniffing) without gathering them to the nest. Normalized latency was calculated using the following formula:

latency = [ (*t*_*1*_ − *t*_*0*_)+ (*t*_*2*_ − *t*_*0*_)+…+ (*t*_*n*_ − *t*_*0*_)] / (n*L)

where *n* = # of pups outside the nest

*t*_*0*_ = start of trial

*t*_*n*_ = time of nth pup gathered

*L* = trial length.

Movement was measured while the animal was performing pup retrieval behavior, using Matlab-based software (MathWorks) [29].

### Injections

Mice were anesthetized with ketamine (100 mg/kg) and xylazine (5 mg/kg) and stabilized in a stereotaxic frame. Lesions in the auditory cortex of CBA/CaJ mice were performed by injection of ibotenic acid (0.5 μL of 10 mg/mL per site; Tocris Bioscience). Control animals were injected with the solvent only (0.9% NaCl solution). Pup retrieval behavior was evaluated 3-5 days later. To knock down MECP2 expression, we injected AAV9-GFP-IRES-Cre (0.3 μL of 4X10e12 molecules/mL per site; UNC Gene Therapy Center) into the auditory cortex of 4 weeks old *MeCP2^flox/flox^* mice. AAV2/7-CMV-EGFP was used as control (both AAV viruses were kind gifts from Dr. Bo Li). Behavior was evaluated 4-6 weeks later. To degrade PNNs, we injected chondroitinase ABC (0.3 μL of 50U/mL per site, in 0.1% BSA/0.9% NaCl solution; Sigma-Aldrich) into the auditory cortex of *MeCP2^het^* and wild type littermate mice. Penicillinase (50U/mL, in 0.1% BSA/0.9% NaCl solution; Sigma-Aldrich) was used as injection control. Pup retrieval behavior was evaluated 3-5 days later. All substances were injected into both auditory cortical hemispheres, 2 sites per hemisphere, at the following coordinates: Bregma = −2.25 and −2.45 mm, ~4 mm lateral and 0.75 mm from the dorsal surface of the brain.

### Immunohistochemistry

Immediately after the behavioral trial on D5, mice were perfused with 4% paraformaldehyde/PBS, and brains were extracted and post-fixed overnight at 4°C. Brains were then treated with 30% sucrose/PBS overnight at room temperature (RT) and microtome sectioned at 50 μm. Free-floating sections were immunostained using standard protocols at RT. Briefly, sections were blocked in 10% normal goat serum and 1% Triton-X for 2-3 hour, and incubated with the following primary antibodies overnight: MECP2 (1:1,000; rabbit; Cell Signaling), PV (1:1,000; mouse; Sigma-Aldrich) and biotin-conjugated Lectin (labels PNNs; 1:500; Sigma-Aldrich). Sections were then incubated with appropriate AlexaFluor dye-conjugated secondary antibodies (1:1,000; Molecular Probes) and mounted in Fluoromount-G (Southern Biotech). To obtain GAD67 staining in soma, three modifications were made according to a previous protocol [30]: 1) no Triton-X or detergent was used in the blocking solution or the antibody diluent; 2) sections were treated with 1% sodium borohydride for 20 minutes prior to blocking, to reduce background; and 3) sections were left in GAD67 antibody (mouse; 1:1,000; Millipore) for 48-60 hours at room temperature. Brains of all uninjected mice were processed together with the mothers at all steps in the process (perfusions, sectioning, immunostaining and imaging with the same settings). Brains of injected mice were processed together with their respective controls at all steps. Brain sections from MECP2 knockdown experiment were further counterstained with a nuclear marker, DAPI.

### Image acquisition and analysis

To analyze percent infection of the auditory cortex by AAV-GFP-Cre or degradation of PNNs by chondroitinase ABC, 4-5 single-plane images per auditory cortical hemisphere from each animal were acquired using Olympus BX43 microscope (4X objective, UPlanFLN) and analyzed using ImageJ (NIH). To calculate percent infection/degradation in each image, the area of the entire auditory cortex was measured based on Allen brain atlas boundaries (Version 1, 2008). Then, the area containing GFP+ cells or reduced PNN expression was measured and divided by the total auditory cortical area. For non-auditory cortical region analysis, cumulative regions included temporal association cortex, ectorhinal cortex and perirhinal cortex. Each correlation data point represents the percent infection/degradation per animal.

To determine the percent of AAV-GFP-Cre infected cells lacking MECP2 expression, 4 confocal images of the auditory cortex (2 images per hemisphere) were acquired using the Zeiss LSM710 confocal microscope (20X objective; 2X zoom) for each AAV-GFP-Cre injected mouse. Using ImageJ (NIH), a region of 100 μm^2^ was used to determine the percent of GFP+ cells that lack MECP2 expression.

For Figure 2F, the amount of MECP2 knockdown was assessed by comparing MECP2 intensity in infected cells (GFP+) and uninfected cells (GFP-) within the same auditory cortical region of each AAV-GFP-Cre injected mouse. 2 confocal images of the auditory cortex (1 image per hemisphere) were acquired using the Zeiss LSM710 confocal microscope (20X objective; 1X zoom) for each mouse. Using ImageJ (NIH) and a region of 150 μm^2^ from each confocal image, the intensity of MECP2 for each GFP+ infected cell was obtained and compared to the intensity of MECP2 in MECP2+ cells that lack GFP (uninfected). Only cells with their entire soma visible in the confocal images were used for the analysis.

**Figure 2:**
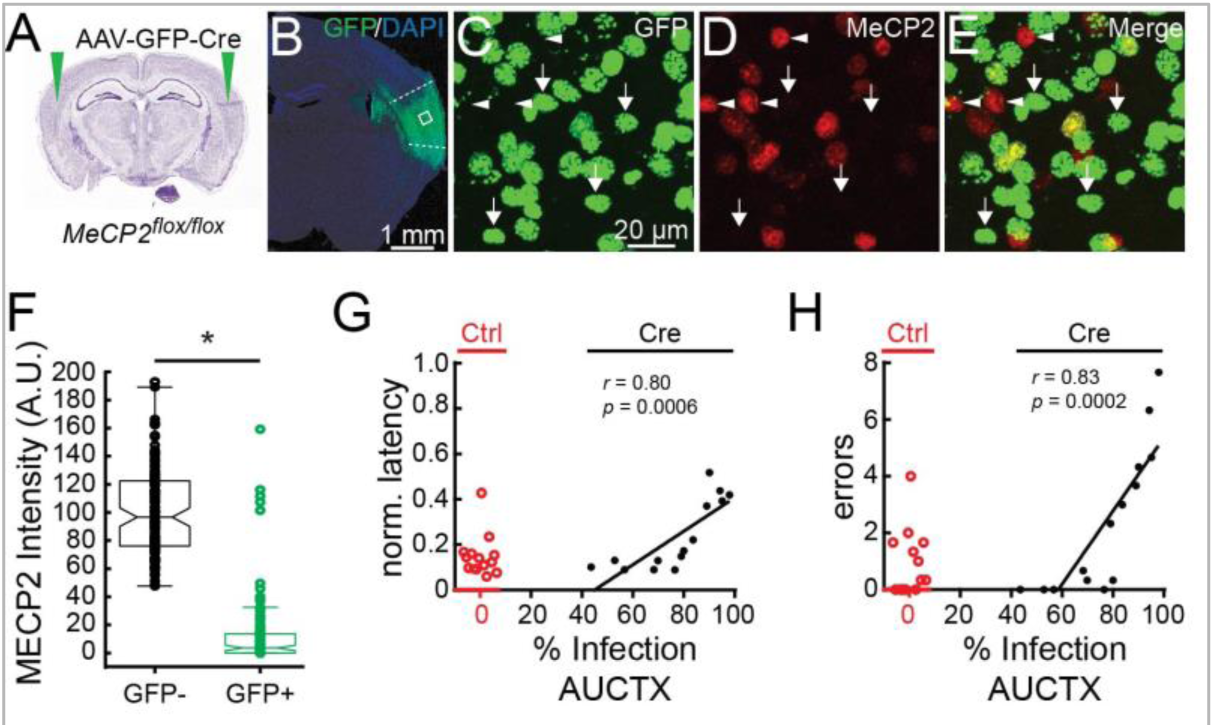
MECP2 expression in the auditory cortex is required for efficient pup retrieval. ***A***, Diagram depicting AAV-GFP-Cre injection into the auditory cortex (green arrows) of female *MeCP2^flox/flox^* mouse. These mice also carried a nuclear localized and Cre-dependent GFP allele (H2B-GFP) that allowed us to directly visualize Cre-positive cells. ***B***, Photomicrograph of a brain section from a *MeCP2^flox/flox^* mouse with AAV-GFP-Cre injection and counterstained with the nuclear marker, DAPI. Dotted lines mark the boundary of auditory cortex. *C-E*, Magnified confocal images of a selected region boxed in B. GFP+ cells (C, green) are negative for MECP2, as confirmed by anti-MECP2 immunostaining (*D and E*, red) (91.2% ± 0.03%; 12 images, 3 mice). Arrows point to GFP+ cells that are MECP2-. Arrowheads point to GFP-cells that are MECP2+, which served as a positive control for MECP2 staining. Scale bar applies to all images. ***F***, Mean MECP2 expression (intensity) in AAV-GFP-Cre infected cells (GFP+) and uninfected cells (GFP-) in the same AAV-GFP-Cre injected animals (n = 119 cells per cell type, 3 animals; *Mann-Whitney, *P < 0.05)*. Cre-infected cells showed significantly reduced MECP2 expression compared to uninfected cells. Boxplot with standard Matlab-generated whiskers are shown. Notches represent 95% confidence interval of median. Each dot overlaid the boxplot represent a cell. ***G, H***, Correlation analysis showed a significant positive relationship between the proportion of auditory cortex expressing GFP-Cre and both gathering latency (G) and number of errors *(H)* (black dots; n = 14 mice; *Pearson’s r)*. Control MeCP2flox/flox mice injected with AAV-GFP alone (Ctrl; red dots) showed normal behavior (n = 14 mice).

To analyze GAD67+ and PV+ soma and PNNs, 2 confocal images from each auditory cortical hemisphere of each animal were acquired using the Zeiss LSM710 confocal microscope (20X objective; 0.6X zoom) and analyzed using the LSM Image Browser. Scans from each channel were collected in the multiple-track mode. Maximum intensity projections of the Z-stacks were obtained using the “Projection” setting in the Zeiss LSM Image Browser. To count high-intensity GAD67+ soma and mature PNNs, the “Contrast” setting in the Browser was set to 100 to threshold weaker signals. GAD67+ soma and mature PNNs were counted manually. Measurement of PV+ cell intensity was performed using Volocity (Perkin Elmer). PV confocal images were first merged. Then, cell identity and intensity were measured using the option “Find 2D nuclei” with “separate touching nuclei = 5 μm” and “reject nuclei of area less than 10 μm^2^.” Results were confirmed manually to exclude non-cell objects and to include any missed PV+ cells. Finally, obtained cell intensities were background subtracted. The experimenter performing the analysis was blinded to all genotypes and conditions. All statistical analysis was performed using Origin Pro (Origin Lab) and Matlab (MathWorks). All graphs were generated using GraphPad Prism (GraphPad Software). Data is represented as mean ± S.E.M.

### In vivo physiology

For awake-state recordings, we anesthetized three *MeCP2^het^* mice with an 80:20 mixture (1.00 mL/kg) of ketamine (100 mg/mL) and xylazine (20 mg/mL) (KX) and stabilized in a stereotaxic frame. A head bar was affixed above the cerebellum with RelyX Luting Cement (3M) and methyl methacrylate-based dental cement (TEETS). For additional support, five machine screws (Amazon Supply) were placed in the skull prior to cement application. After one day of recovery, mice were anesthetized with isoflurane (Fluriso; Vet One) and small craniotomies were made to expose the left hemisphere of auditory cortex. Mice were then head-fixed via the attached head bar over a Styrofoam ball that was suspended above the air table. The Styrofoam ball allowed mice to walk and run in one dimension (forward-reverse).

For anesthetized-state recordings, we anesthetized a *MeCP2^het^* mouse with KX and the left auditory cortex was exposed with small craniotomy. Animal was placed in a warming tube and kept anesthetized with injection of 10 uL of ketamine as needed throughout the recording session.

Stimuli were presented via ED1 Electrostatic Speaker Driver and ES1 Electrostatic Speaker (TDT), in a sound attenuation chamber (Industrial Acoustics) at 65 dB SPL RMS measured at the animal’s head. Stimuli consisted of broadband noise, four logarithmically-spaced tones ranging between 4 and 32 kHz, and ultrasound noise bandpassed between 40 and 60 kHz.

Single units were blindly recorded by *in vivo* loose-patch technique using borosilicate glass micropipettes (7-30 MΩ) tip-filled with intracellular solution (mM: 125 potassium gluconate, 10 potassium chloride, 2 magnesium chloride and 10 HEPES, pH 7.2). Spike activity was recorded using BA-03X bridge amplifier (npi Electronic Instruments), low-pass filtered at 3 kHz and digitized at 10 kHz, and acquired using Spike2 (Cambridge Electronic Design). Data were analyzed using Spike2 and Matlab.

To assess statistical significance of responses to individual odors, we used a bootstrap procedure as follows. If *n* trials were collected with the response window length *t*, then a distribution was created by sampling *n* length *t* windows from the full spike record 10,000 times and taking the mean deviation of each window from the spike rate measured in the prior 2 s. Responses that were in the top or bottom 2.5% of this distribution were deemed significantly excitatory or inhibitory, respectively.

## RESULTS

### Pup retrieval is a learned behavior that requires the auditory cortex

To assess the efficacy of cortical plasticity underlying pup-retrieval learning, we devised an assay for gathering behavior in nulliparous surrogates (*Sur*). We chose to examine cortical plasticity underlying the acquisition of gathering behavior in surrogates to eliminate the influence of pregnancy. Our intent was not to study maternal behavior *per se* or plasticity in mothers, but to use this assay to study the function of MECP2 in adult experience-dependent plasticity in surrogates at the neural circuit and behavioral levels. Assaying the effects of heterozygous deletion of *MeCP2* on gathering behavior presents several advantages. First, the vast majority of RTT patients are females heterozygous for mutations of *MeCP2* who exhibit mosaic expression of the wild type protein. Thus, female *MeCP2^het^* [31] are a particularly appropriate model of RTT. Second, we can directly relate a natural, learned adult behavior to specific, experience-dependent changes in the underlying neural circuitry. Third, we can observe effects on adult learning and plasticity that are distinct from developmentally programmed events in the surrogates by studying a window of heightened plasticity that is triggered by exposure to a mother and her pups.

Two 7 - 10 weeks old matched female littermates (*Sur*) were co-housed with a first time mother and her pups from late pregnancy through the fifth day following birth (D5) (Figure 1A). Surrogates were virgins with no prior exposure to pups. All three adults (the mother and both surrogates) were subjected to a retrieval assay (see Materials and Methods) on D0 (day of birth), D3 and D5.

**Figure 1:**
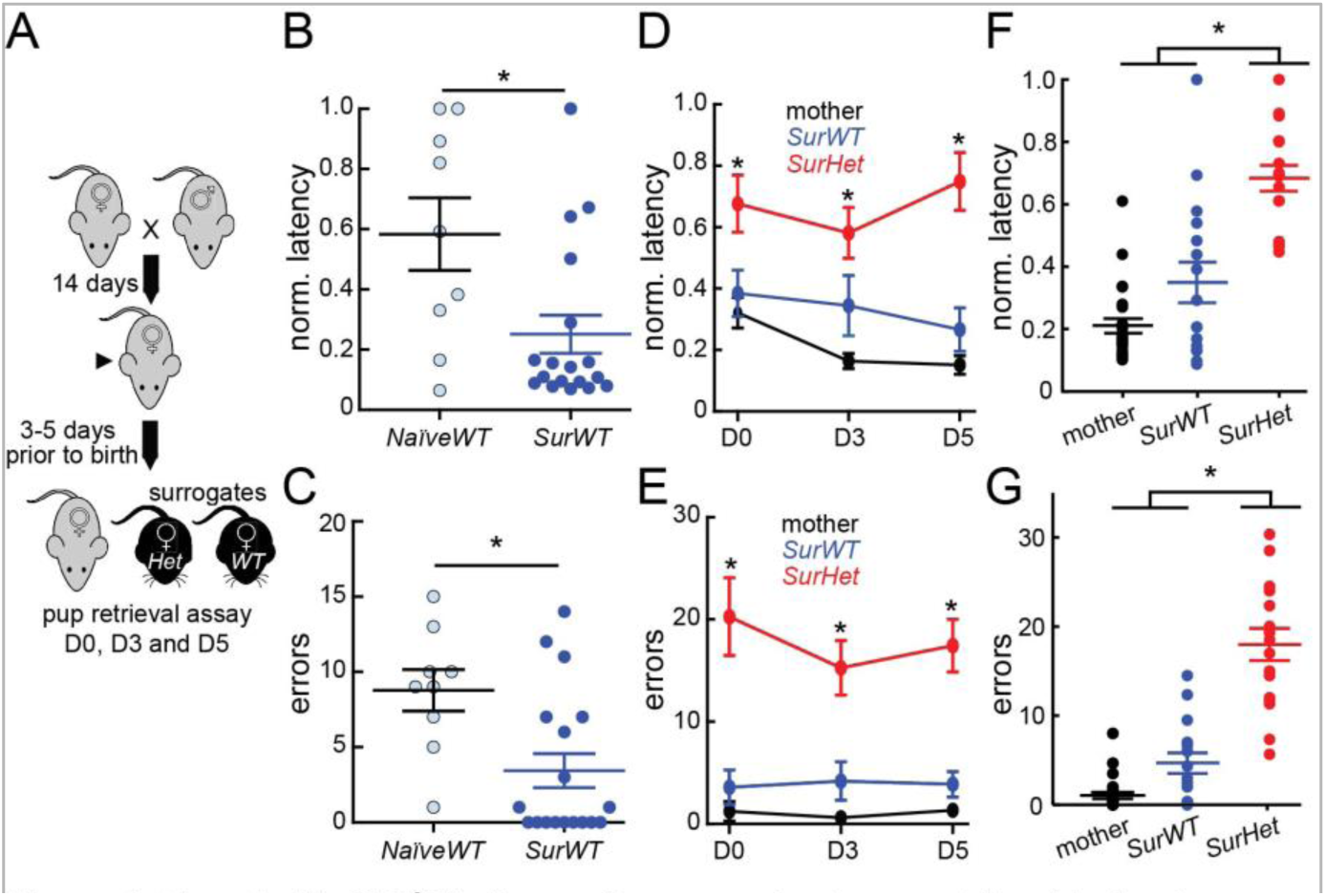
Female *MeCP2^het^* mice perform poorly at pup retrieval behavior. **A**, Schematic of behavioral paradigm. Virgin *MeCP2^het^ (Het)* and wild type littermates (WT) mice were co-housed with a pregnant female prior to birth of pups. Surrogates were tested on the pup retrieval task on days 0, 3 and 5 after birth. ***B, C***, Surrogate WTs *(SurWT)* tested on D5 (dark blue; n = 18 mice) showed significant improvements on a normalized measure of latency to gather (B) and reduced number of gathering errors *(C)*, compared to pup-naive mice (grey; n = 9 mice) *(Mann-Whitney U Test, *P = 0.027). **D, E***, Mean performance at DO, D3 and D5 for mothers, SurWT and surrogate *MeCP2^het^ (SurHet)* as measured by normalized latency *(D)* and errors *(E). SurHet* showed consistently poorer pup retrieval performance than mothers and *SurWT* in all three sessions (n = 13-24 mice; *Kruskal-Wallis with Bonferroni correction, *P < 0.05). **F, G***, Mean performance of normalized latency (F) and errors (G) averaged over all three sessions (n = 13-24 mice; *Kruskal-Wallis with Bonferroni correction, *P < 0.05). SurHet* had significantly longer latency and made more errors compared to mothers and *SurWT*. Mean ± S.E.M. are shown in line.

We confirmed the experience-dependent nature of gathering behavior by comparing performance of maternally-naïve WT (*NaïveWT*) females with that of *SurWT* on D5. Performance was assessed by computing a normalized measure of latency (see Materials and Methods) and by counting the number of gathering errors (instances of interacting with a pup and failing to gather it to the nest). *SurWT* performed significantly better than *NaïveWT* by both measures (Video 1) (Figure 1B, C) (*NaïveWT*: n = 9 mice; *SurWT*: n = 18 mice; *Mann-Whitney U Test, *P = 0.027)*, presumably reflecting maternal experience-dependent plasticity.

Several lines of evidence suggest that auditory cortical responses to ultrasonic pup vocalization (USVs) facilitate performance of pup gathering behavior [16-18] [32]. We confirmed this by making bilateral excitotoxic (ibotenic acid) lesions of the auditory cortex in wild type mice. Compared to saline-injected mice, mice with lesions exhibited significantly longer latency (Saline: 0.20 ± 0.034, n = 6 mice; Lesion: 0.66 ± 0.033, n = 6 mice; *Mann-Whitney: P = 0.0022*) and made more errors (Saline: 1.33 ± 0.95, n = 6 mice; Lesion: 6.64 ± 0.91, n = 6 mice; *Mann-Whitney: P=0.015)*.

### MECP2 is required for efficient pup retrieval behavior

Next, we compared the gathering performance of *SurHet* with that of mothers and *SurWT. SurWT* retrieved pups to the nest with efficiency (as measured by latency in Figure 1D, F) and accuracy (as measured by errors in Figure 1E, G) that were indistinguishable from the mother. *SurHet* exhibited dramatic impairment in gathering behavior, retrieving pups with significantly longer latency and more errors when compared to the *SurWT* or mothers (Figure 1D-G). Moreover, this behavior does not improve with subsequent testing on D3 and D5 (Figure1D, E) (n = 13-24 mice; *Kruskal-Wallis with Bonferroni correction, *P < 0.05*), demonstrating that MECP2 is required for successful acquisition of this behavior.

In these experiments, we used a germline *MeCP2* knockout that affects MECP2 expression throughout the brain. Therefore the poor behavior of *SurHet* could in principle be due to motor deficits or deafness. We found no evidence of impaired movement by *SurHet* during testing. There was no significant difference in movement during behavioral trials between the genotypes (*SurWT*: 2059 ± 216.5 significant motion pixels (SMP), n = 8 mice; *SurHet*: 2139 ± 259.9 SMP, n = 8 mice; *Mann-Whitney: P = 0.78*), consistent with previous findings that *MeCP2^het^* lack robust motor impairments [33]. Furthermore, neuronal responses we observed in the auditory cortex clearly demonstrate that the ear and early auditory system are functional. Neurons in the auditory cortex of *NaïveHe*t exhibited widespread and robust responses to auditory stimuli. Among all neurons recorded in three *NaiveHet* (see Materials and Methods), 27/28 cells responded significantly to at least one of six tone and noise stimuli. The mean number of stimuli that evoked a significant response in the 28 neurons was 2.79 ± 1.4 stimuli. Therefore, the impaired gathering behavior in *SurHet* is not caused by deafness or insensitivity of the early auditory system.

### MECP2 expression in the adult auditory cortex is required for efficient pup retrieval behavior

Measuring behavioral effects in germline mutants leaves open the possibility of a requirement for MECP2 in early postnatal development and/or in other brain regions. Therefore, we used a conditional deletion approach to specifically remove MECP2 in the auditory cortex by injecting AAV-GFP-Cre (adeno-associated virus expressing CRE recombinase) in 4-week old *MeCP2^flox/flox^* mice [34] (Figure 2A). Histological analysis of sections from *SurMeCP2^flox/flox^* five weeks Histological analysis of sections from *SurMeCP2^flox/flox^* five weeks after injection with AAV-Cre showed >91% of GFP+ nuclei (12 images, 3 animals) (see Materials and Methods) were devoid of MECP2 (Figure 2B-F). We counted GFP (-) and GFP (+) cells to determine the extent of MECP2 knock-down in the GFP (+) cells and found significant reduction of MECP2 expression in the GFP (+) cells of the auditory cortex (Figure 2F; n = 119 cells per cell type, 3 animals; *Mann-Whitney, *P < 0.05*).

*MeCP2^flox/flox^* mice injected with AAV-GFP alone (control), consistently showed strong gathering performance (Figure 2 G, H) (n = 14 animals). In contrast, *MeCP2^flox/flox^* mice injected with AAV-Cre exhibited variable gathering behavior that depended on the proportion of auditory cortex affected by the injection. The degree of impairment for an individual mouse was significantly positively correlated with the percentage of the auditory cortices encompassed by the virus injection site (Figure 2G, H) (Latency: *r* = 0.80, *p* = 0.0006; n = 14 animals; Errors: *r* = 0.83, *p* = 0.0002; n = 14 animals; *Pearson’s r*). No positive correlation between injection area and behavioral performance was found with regions surrounding the auditory cortex (Latency: *r* = 0.40, *p* = 0.16; Errors: *r* = 0.25, *p* = 0.40; n = 14 animals; *Pearson’s r*). Taken together, these findings demonstrate that normal MECP2 expression specifically in the auditory cortex of mature females is required for proficient gathering behavior.

### SurHet exhibit dysregulated plasticity of GABAergic interneurons in the auditory cortex

The regional requirement for MECP2 led us to examine maternal experience-dependent molecular events in the auditory cortex. Recent data on the neurophysiological correlates of maternal learning suggest that there are changes in inhibitory responses of vocalizations in the auditory cortex of mothers and surrogates [17, 35]. There is also evidence that inhibitory networks are particularly vulnerable to *MeCP2* mutation [36-38]. For these reasons, we focused our attention on experience-dependent dynamics of molecular markers associated with inhibitory circuits.

We used immunostaining of brain sections from the auditory cortex of surrogates and naïve females to examine experience-induced molecular events in inhibitory networks of *MeCP2^het^* and wild-type littermates. Expression of GAD67, the key rate-limiting enzyme for GABA synthesis was significantly increased five days after initiation of maternal experience in mutant and wild type mice (Figure 3A, B) (n = 12-32 images, 4-8 mice; *ANOVA: Tukey’s post-hoc test, ^*^P < 0.05*). For both surrogate genotypes, expression returned to baseline by the time the pups were weaned (D21) (Figure 3A, B). This suggests that maternal experience triggers transient experience-dependent molecular changes in inhibitory neurons in the auditory cortex of surrogate mice.

**Figure 3:**
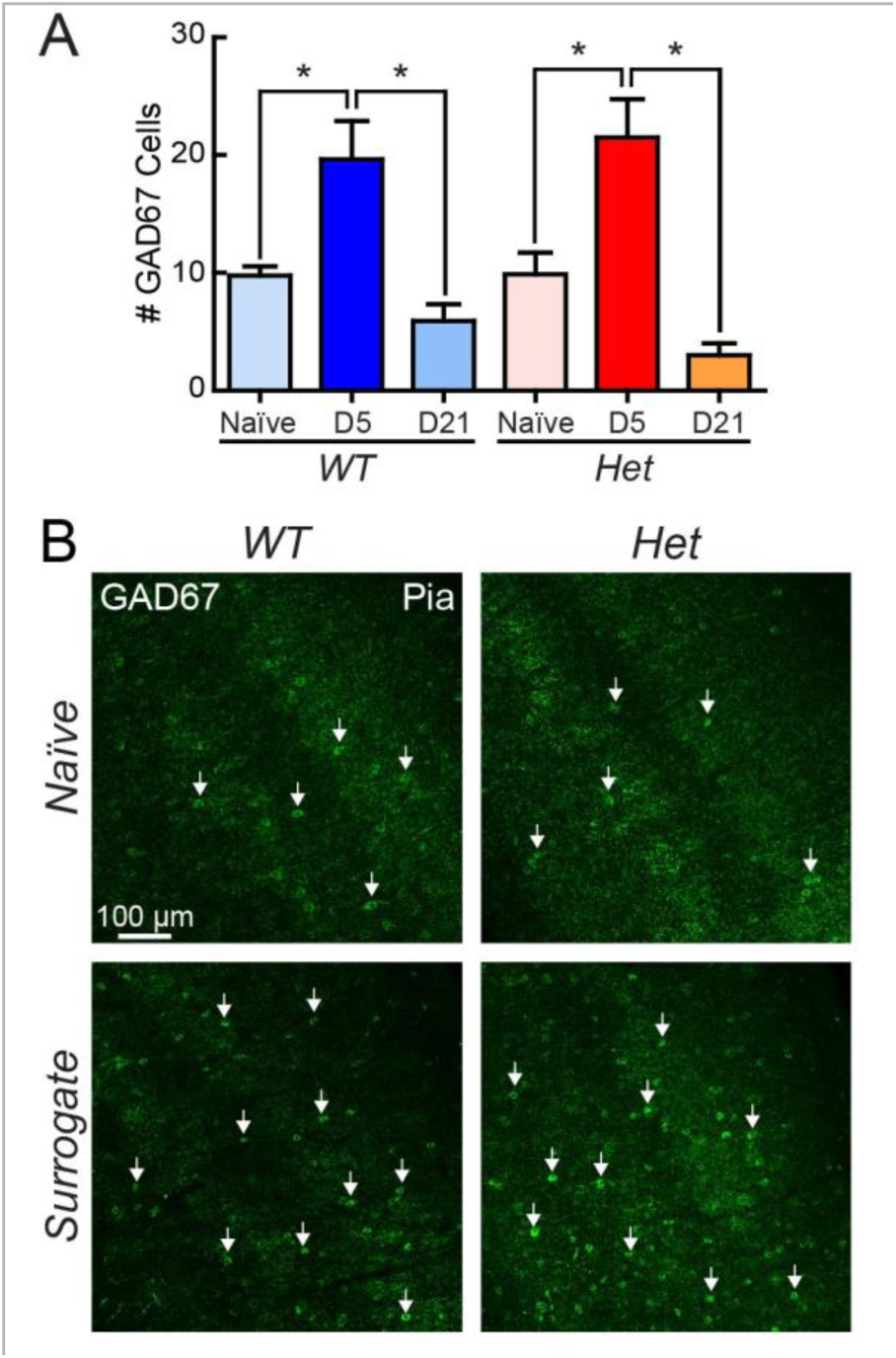
Maternal experience transiently enhances GAD67 expression level in the auditory cortex of wild-type and MeCP2het mice. ***A***, The number of high-intensity GAD67 cells was significantly increased in both *SurWT* (dark blue) and *SurHet* (red) at D5, and returned to naive levels at D21 (n = 12-32 images, 4-8 mice; *ANOVA: Tukey’s post-hoc test, *P < 0.05)*. Bar graphs represent mean + S.E.M. B, Representative confocal images taken from the auditory cortex of a *NaïveWT* and *NaïveHet* (top row) and *SurWT* and *SurHet* at D5 (bottom row). Arrows point to high-intensity GAD67 cells. Scale bar applies to all images.

In *SurHet* only, we observed transient increases in additional markers of inhibitory networks that are often associated with suppressing plasticity. For example, recent work has linked high parvalbumin (PV)-expressing inhibitory networks to reduced capacity for adult learning and plasticity [25] and the closure of developmental critical periods [21, 24, 39]. We detected a maternal experience-induced shift in the intensity distribution of PV immunofluorescence in *SurHet* but not *SurWT* (Figure 4A, B, D). The intensity distribution for *SurHet* was fit with a mixture of two Gaussians to define high and low PV-expressing populations. The proportion of high PV-expressing neurons was significantly greater in *SurHet* than in any other group (Figure 4B) (n = 19-20 images, 5 mice for each group; *ANOVA: Tukey’s post-hoc test, ^#^P < 0.05* compared to all other groups).

**Figure 4:**
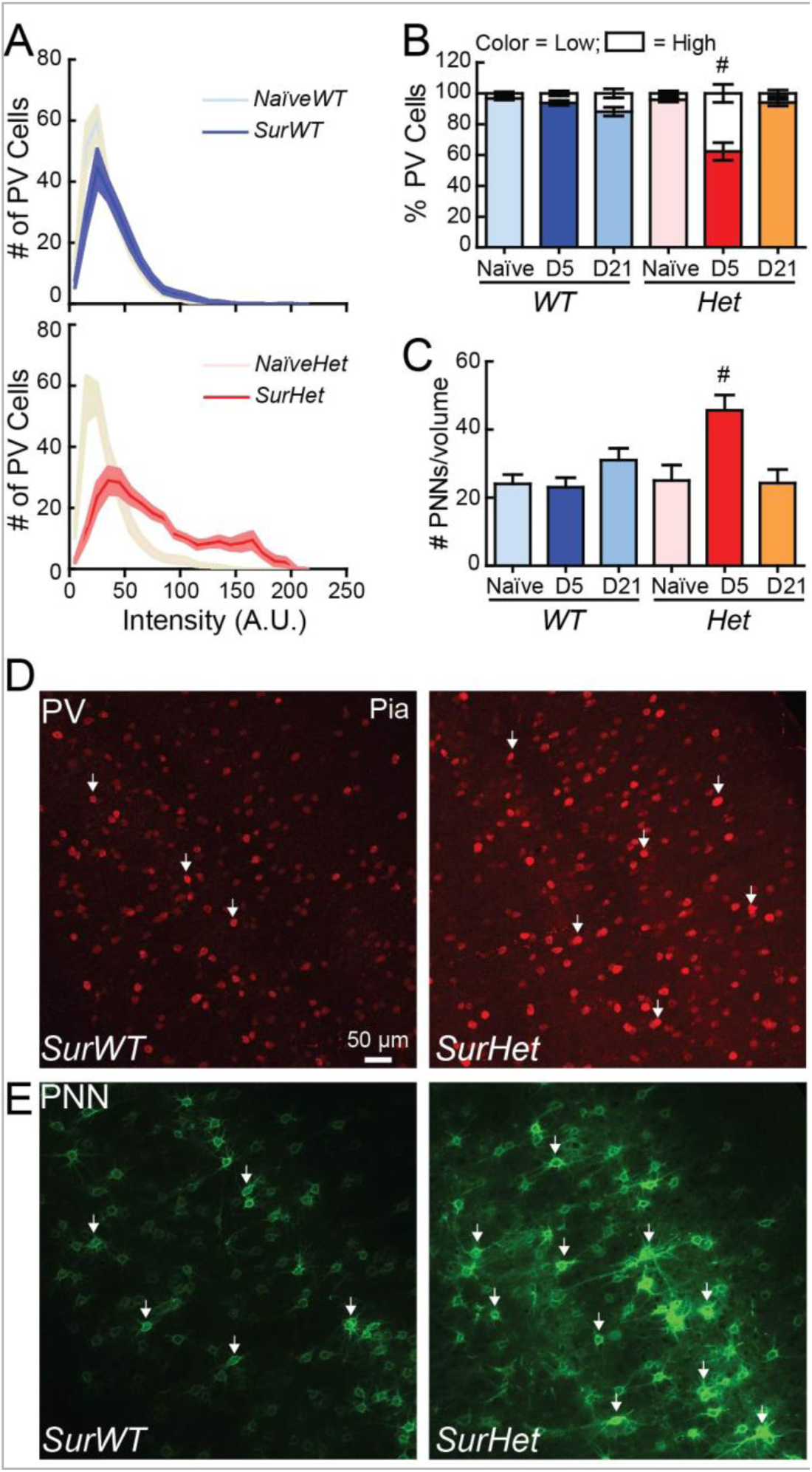
Female *MeCP2^het^* mice exhibit abnormal maternal experience-induced changes to inhibitory networks in the auditory cortex. ***A***, Histograms showing the mean distribution of PV cell intensity in adult surrogates 5 days after pup exposure (D5). Top panel, Distribution of PV cell intensity is similar between *SurWT* (dark blue) and *NaïveWT* (light blue). Bottom panel, There is a shift in the distribution toward elevated PV expression in *SurHet* (red) compared to *NaïveHet* (pink) (n = 19-20 images, 5 mice for each group). The solid line and shaded region represent mean ± S.E.M. respectively, in both panels. ***B***, The shift reflects a significant transient increase in high-PV expressing cells at D5 (red) that returned to baseline at D21 (orange) in *SurHet* (*ANOVA: Tukey’s post-hoc test, ^#^P < 0.05* compared to all other groups). ***C***, The number of high-intensity perineuronal nets (PNNs) was significantly increased only in *SurHet* at D5 (red) (n = 12 38 images, 3-9 mice; *ANOVA: Tukey’s post-hoc test, ^#^P<0.05* compared to all other groups), and returned to baseline at D21 (orange). ***D, E***, Representative confocal images taken from the auditory cortex of a *SurWT* and *SurHet* showing relative expression of PV *(D)* and PNN *(E)*. Arrows point to high-intensity PV and PNN in both genotypes. Scale bar applies to all images.

Mature neural circuits are often stabilized by perineuronal nets (PNNs), which are composed of extracellular matrix proteins such as chondroitin sulfate proteoglycans (CSPGs) [40], and mainly surround PV+ GABAergic interneurons in the cortex [41]. We observed a dramatic experience-dependent increase in the number of high-intensity PNNs in *SurHet* but not in *SurWT* (Figure 4C, E) (n = 12-38 images, 3-9 animals; *ANOVA: Tukey’s post-hoc test, ^#^P < 0.05* compared to all other groups). Importantly, both PV and PNNs returned to baseline levels in surrogates by weaning age of the pups (D21) (Figure 4B, C).

Thus, maternal experience triggers temporally-restricted changes to inhibitory circuits in *SurHet*, but there are additional changes not seen in *SurWT*, including elevated PV and PNN expression. We have separately observed elevated PV and PNN expression and altered plasticity in the visual cortex of *MeCP2-null* males during the visual critical period [42]. Similar changes may act to limit network plasticity after maternal experience. Moreover, the reversion to baseline levels following weaning indicates that pathological features of the plasticity are temporally limited and suggests that certain aspects of *MeCP2^het^* pathology are only revealed during appropriate experiences within that window.

### Rescue of cellular and behavioral phenotypes of SurHet by Gad1 genetic manipulation

GAD67 is an activity-regulated, rate-limiting enzyme that synthesizes the cortical inhibitory neurotransmitter GABA. GAD67 expression levels also correlate highly with PV levels [25] and regulate PV neuron maturation [43]. Several recent studies suggest that mice that are heterozygous for loss of the GAD67 gene (*Gad1*) exhibit lower levels of PV expression (Uchida, 2014) [44]. We have separately observed that lowering GAD67 levels in the *MeCP2-null* male mice normalized expression of PV and PNN in the developing visual cortex [42]. We therefore speculated that genetically manipulating GAD67 expression (*Gad1^het^*) in *MeCP2^het^* might result in normalization of PV network associated markers in the adult auditory cortex. To test this idea, we crossed germline *Gad1^het^* mice into the *MeCP2^het^* background and examined the effects on maternal experience-dependent changes in PV and PNNs.

As expected, naïve WT and *MeCP2^het^* carrying the *Gad1^het^* allele (*Gad1^het^* and *NaïveHet;Gad1^het^*, respectively) showed half the GAD67 expression seen in WT and *MeCP2^het^* (*NaïveWT*: 11.5 ± 1.5 cells, *NaïveHet*: 9.9 ± 1.8 cells, *NaïveHetGad1^het^*: 4.9 ± 1.0 cells, *NaiveGad1^het^*: 4.4 ± 1.5 cells; n = 20-32 images, 5-8 animals; *T-test: P < 0.05 NaïveHetGad1^het^* compared to *NaïveWT* and *NaïveHet; T-test: P < 0.05 NaïveGad1^het^ compared to NaïveWT and NaïveHet*). In contrast to *SurHet*, *SurHet*;*Gad1^het^* exhibited a correction in the maternal experience-dependent increase in PV expression levels (Figure 5A, B) and had a significantly lower proportion of high-intensity PV+ cells (Figure 5B) (n = 16-20 images, 4-5 mice; *ANOVA: Tukey’s post-hoc test, ^*^P = 0.02*). We also saw significantly fewer PNNs in the double mutants (Figure 5D) (n = 17-38 images, 4-9 animals; *ANOVA: Tukey’s post-hoc test, *P = 0.01*). *NaïveGad1^het^* exhibited a significantly elevated number of high-intensity PV+ cells, compared to *NaïveWT* (Figure 5C), likely due to compensatory effects of long-term genetic reduction of GAD67. Interestingly, this increase was not seen after maternal experience (Figure 5C), returning to the appropriate activity-dependent expression of PV in the wild type background (n = 16-20 images, 4-5 animals; *ANOVA: Tukey’s post-hoc test, *P < 0.05)*. There was no change in PNNs in this genotype, before or after maternal experience (Figure 5E) (n = 16-28 images, 4-7 animals; *ANOVA: Tukey’s post-hoc test, P > 0.05*). These data indicate that manipulating GAD67 in the *MeCP2*-deficient background is effective in ameliorating pathological features of maternal experience-dependent auditory cortical plasticity in *SurHet*.

**Figure 5:**
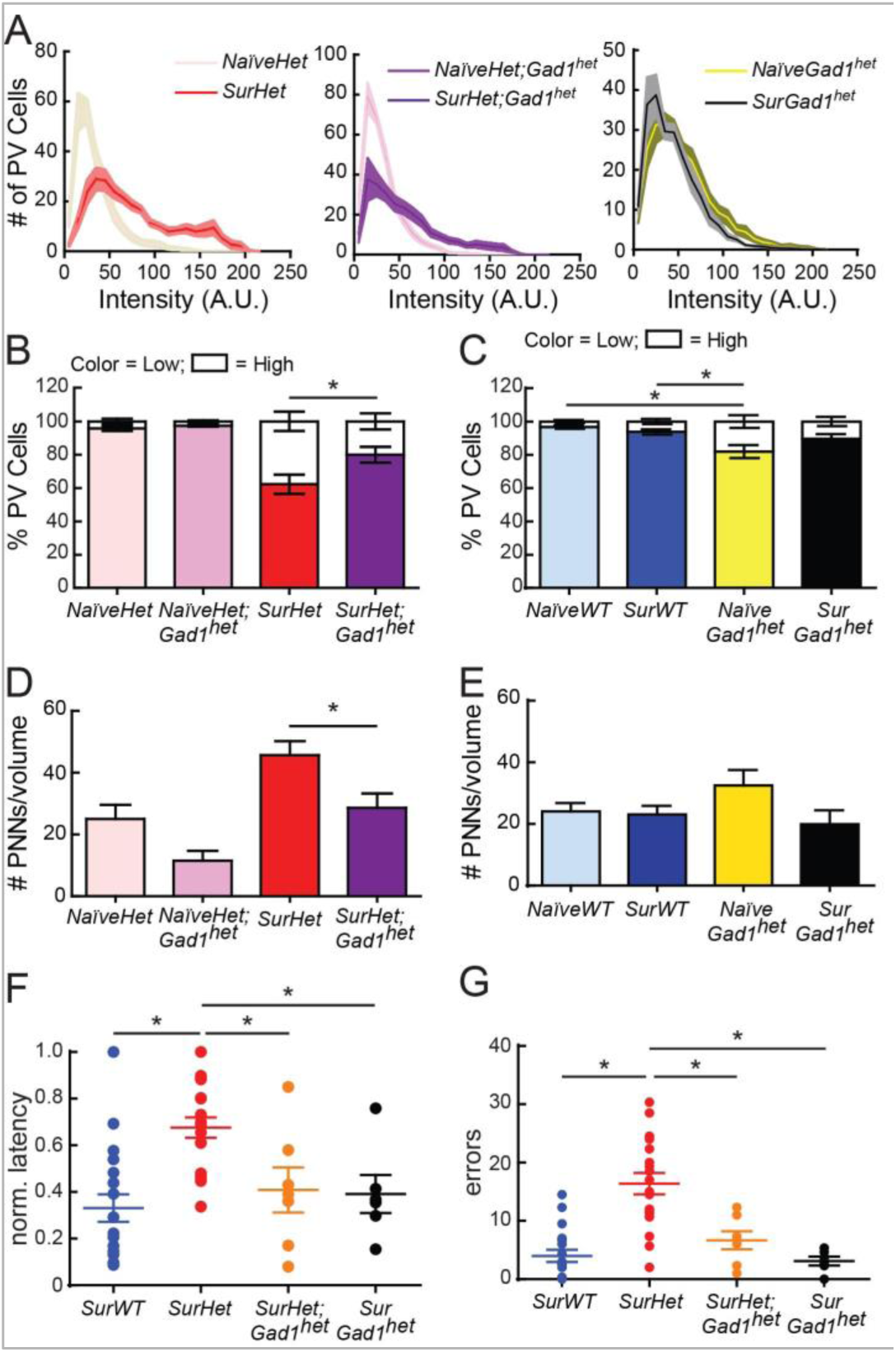
Genetic manipulation of the GABA synthesizing enzyme *Gad1* rescues cellular and behavioral phenotypes in *MeCP2^het^*. ***A***, Histograms showing the mean distribution of PV cell intensity comparing *SurHet* (left, red), *SurHet;Gad1^het^* (middle, purple) and *SurGad1^het^* (right, black) at D5 to their respective naive genotypes. *SurHet;Gad1^het^* showed a smaller shift toward elevated PV expression after maternal experience (n = 16-20 images, 4-5 mice). The solid line and shaded region represent mean ± S.E.M., respectively. ***B**, SurHet;Gad1^het^* (purple) showed a significant decrease in the high-intensity PV population compared to *SurHet* (red) at D5 *(ANOVA: Tukey’s post-hoc test, *P = 0.02). **C**, NaïveGad1^het^* showed significantly more high-intensity PV cells compared to *NaïveWT* and *SurWT*. Upon maternal experience, PV population of *SurGad1^het^* shifted to WT PV expression level. (n = 16-20 images, 4-5 animals; *ANOVA: Tukey’s post-hoc test, *P < 0.05). **D***, At D5, high-intensity PNN counts were significantly reduced in *SurHet;Gad1^het^* (purple), compared to *SurHet* (n = 17-38 images, 4-9 mice; *ANOVA: Tukey’s post-hoc test, *P = 0.01). **E***, High-intensity PNN counts were not significantly different between *Gad1^het^* and wild type mice, before and 5 days after maternal experience (n = 16-28 images, 4-7 animals; *ANOVA: Tukey’s post-hoc test, P > 0.05). **F, G***, Pup retrieval behavior is significantly improved in *SurHet;Gad1^het^* (purple) (n = 8) as measured by normalized latency (F) and errors (G) averaged across three sessions *(SurWT:* n = 18 mice; *SurHet:* n = 18 mice; *SurGad1^het^:* n = 7 mice. *ANOVA: Tukey’s post-hoc test*, \**P* < *0.05*). Mean ± S.E.M. are shown.

We next assessed whether the corrective effect of GAD67 reduction on inhibitory markers in *SurHet* reinstated learning. Remarkably, *SurHet*;*Gad1^het^* exhibited significant decreases in latency (Figure 5F) and number of errors (Figure 5G) (*SurWT*: n = 18 mice; *SurHet*: n = 18 mice; *ANOVA: Tukey’s post-hoc test, *P < 0.05*) when compared to *SurHet*. In fact, the gathering performance of *SurHet*;*Gad1^het^* was indistinguishable from that of *SurWT* or *SurGad1^het^* (Figure 5F, G). These results show that manipulating GABAergic neurons in the *MeCP2*-deficient background alleviates learning deficits, potentially through effects on levels of PV and PNNs.

### Suppressing PNN formation in auditory cortex of SurHet improves maternal learning

PNNs are known to act as barriers to structural plasticity [23, 24]. Thus, relief from the excessive formation of PNNs in *SurHet*;*Gad1^het^* could be a critical factor allowing efficient pup gathering. We speculated that suppressing PNN formation selectively in the auditory cortex just before maternal experience is sufficient to improve behavioral performance of *SurHet*. We therefore made bilateral auditory cortical injections of chondroitinase ABC (ChABC), which dissolves and suppresses the formation of PNNs [24], thereby allowing for the formation of new synaptic contacts [40].

Two sites of injection were made into each hemisphere one to three days prior to initiating assessment of retrieval performance (see Materials and Methods). Injection of ChABC into the auditory cortex of *Het* and *WT* significantly reduced high-intensity PNN counts compared to their respective controls: penicillinase [45] injected mice (Figure 6A-C) (*Pen-Het*: n = 31 images, 8 mice; *ChABC-Het*: n = 24 images, 6 mice; *Mann Whitney, ^*^P = 0.0003; Pen-WT* and *ChABC-WT*: n = 32 images, 8 animals per condition; *Mann Whitney: *P < 0.0001*).

**Figure 6:**
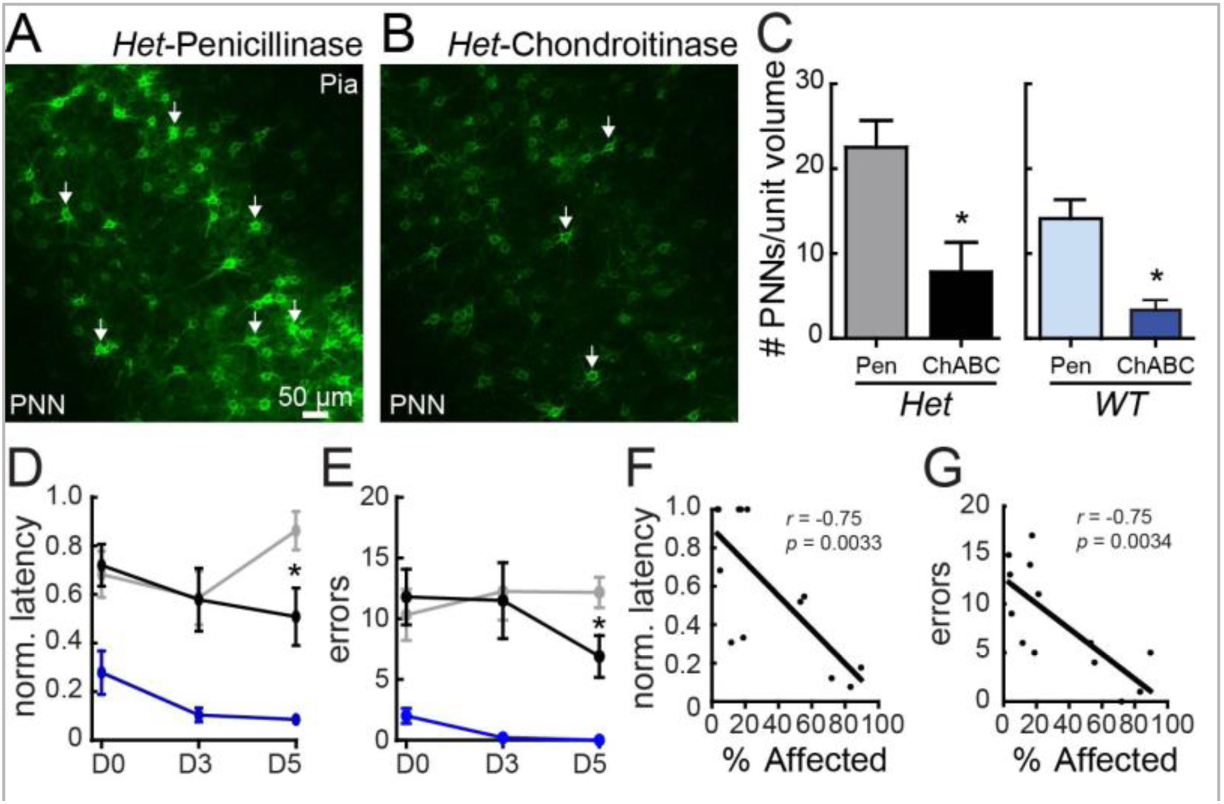
Pharmacological suppression of PNN formation in the auditory cortex restores wild-type behavior in *MeCP2^het^*. ***A, B***, Sample confocal images taken at D5, from the auditory cortex of *SurHet* that received injections of either a control enzyme *(A*, penicillinase) or an enzyme that dissolves PNNs (B, chondroitinase ABC). Arrows indicate high-intensity PNNs. ***C***, At D5, chondroitinase ABC (ChABC) significantly dissolved PNNs in the injected brains of *Het* and *WT*, compared to their respective penicillinase (Pen)-injected genotypes *(Pen-Het* n = 31 images, 8 animals, *ChABC-Het:* n = 24 images, 6 animals, *Mann-Whitney, *P* = *0.0003; Pen-WT* and *ChABC-WT:* n = 32, 8 animals per condition, *Mann-Whitney, *P < 0.0001). **D, E***, Pup retrieval behavior improved significantly on D5 in *SurHet* injected with chondroitinase ABC (black), as measured by normalized latency (D) and errors (E), compared to penicillinase-injected *SurHet* (grey) (Penicillinase: n = 12 mice; ChABC: n = 13 mice; *ANOVA: Tukey’s post-hoc test, *P < 0.05)*. No significant differences in latency and error were observed between ChABC-injected and penicillinase-injected WT except at D3 *(Mann Whitney, P* = *0.048)*. For simpler graphic presentation, only ChABC-injected WT data are shown in blue *(Pen-WT:* n = 7 animals; *ChABC-WT:* n = 5 animals). Mean ± S.E.M. are shown. ***F, G***, Correlation analysis showed a significant negative relationship between the proportion of auditory cortex encompassed by chondroitinase ABC injection for both latency (F) and number of errors (G) *(Pearson’s r)*.

*SurHet* mice that received bilateral injections of ChABC in the auditory cortex showed significantly improved gathering performance of D5 pups. ChABC-injected *SurHet* retrieved pups with lower latency (Figure 6D) and fewer errors (Figure 6E) compared to *SurHet* injected with the control enzyme, penicillinase [24] (Het-Pen: grey line, n = 12 mice; ChABC-Het: black line, n = 13 mice; *Mann Whitney, ^*^P < 0.05*). ChABC-injected *WT* performed similarly to the penicillinase-injected *WT*, with a small significant decrease in latency at Day 3 (Figure 6D, E) (For clarity, only ChABC-WT data are shown in blue line, n = 5 animals; Pen-WT: normalized latency – D0 = 0.43 ± 0.13, D3 = 0.29 ± 0.12, D5 = 0.19 ± 0.07, n = 7 animals; At D3: *Mann Whitney, P = 0.048*; Pen-WT: errors – D0 = 2.3 ± 1.4, D3 = 1.6 ± 0.78, D5 = 1.57 ± 0.81, n = 7 animals; *Mann Whitney, P > 0.05*).

Due to technical variability, not all injections covered the entire auditory cortex. Hence, we correlated the percentage of the region affected by the injection with gathering performance. In *SurHet*, the proportion of auditory cortex bilaterally encompassed by the injection site was significantly negatively correlated with latency (Figure 6F) (n = 13 animals, *r = −0.75, p = 0.0033, Pearson’s r*) and number of errors (Figure 6G) (n = 13 animals, *r = −0.75, p = 0.0034, Pearson’s r*) exhibited on D5 pups. Interestingly, this relationship did not emerge until day 5 of maternal experience. Therefore, increased PNNs in *SurHet* inhibit auditory cortical plasticity required for rapid and accurate pup gathering.

## DISCUSSION

A key challenge for understanding the pathogenesis of RTT and neuropsychiatric disorders in general is to identify the associated molecular/cellular changes and trace the resulting circuit alterations that underlie behavior deficits. It is also critical to differentiate between impairment of developmental programs and effects on experience-dependent neural plasticity. Here we take advantage of a robust natural behavior in female mice that relies on a known cortical region, and we make a clear and direct link between molecular events in that region and behavior. Our data identify a specific critical role for MECP2 in experience-dependent plasticity of cortical inhibitory networks in adults.

Most previous studies in mouse models of RTT were conducted in *MeCP2*-null male mouse models because they exhibit earlier and more severe phenotypes in many assays. Therefore, with the exception of a few studies [33, 46, 47], the molecular, circuit and behavioral defects in *MeCP2^hets^* are unknown. We found a robust behavioral phenotype in these mice that reveals a dramatic impairment of adult experience-dependent plasticity. Importantly, *MeCP2^hets^* are genetically a more accurate model of human RTT, so our data may likely reflect a critical component of the human disorder as well. Based on our results, we conclude that dysregulated auditory processing, likely due to impaired plasticity, leads directly to altered behavior. Notably, we also showed that when normal plasticity is restored, even acutely in adulthood, behavior improves.

### MeCP2 mutation interferes with experience-dependent plasticity and behavior in adults

Emerging evidence indicates that appropriate expression and function of MECP2 is required in adulthood for normal plasticity and behavior [8, 9]. Remarkably, restoring normal MECP2 expression in adulthood improves symptoms [46, 48]. These observations have several implications. First, they indicate that some cellular functions of MECP2 are involved in maintenance and adult plasticity of neural circuitry as opposed to only its development. Second, they raise the possibility that there may be a benefit to intervention at all stages of MECP2 deficiency. Nevertheless, the specific mechanisms by which *MeCP2* mutations impair adult function need to be elucidated.

Our data demonstrate that heterozygous mutations in *MeCP2* (*MeCP2^+/−^*) interfere with auditory cortical plasticity that occurs in adult mice during initial maternal experience. Mothers and wild type virgin surrogates achieve proficiency in pup retrieval behavior by an experience-dependent learning process [16, 19, 20, 35, 49, 50], that is correlated with physiological plasticity in the auditory cortex [16-18, 51]. We used gathering behavior as a ‘read-out’ to assay defects in this sensory plasticity. Our results clearly show that *MeCP2^hets^* have markedly impaired ability to learn appropriate gathering responses to pup calls. This interference is in large part due to a specific requirement for MECP2 in adult auditory cortex. Deletion of MECP2 in adult mice selectively in the auditory cortex also produced inefficient retrieval.

### Maternal experience triggers an episode of inhibitory plasticity that is disrupted by MeCP2 mutation

We find evidence of dysregulated cortical inhibitory networks during maternal experience in *MeCP2^het^*. This is consistent with increasing evidence that dysfunction of GABA signaling is associated with autism disorders and RTT [36-38, 52]. Importantly, disruption of MECP2 in GABAergic neurons recapitulates multiple aspects of RTT including repetitive behaviors and early lethality [37], though the pathogenic mechanisms remain unclear.

Our data suggest that an important aspect of the pathology associated with heterozygous *MeCP2* mutations is impaired plasticity of cortical inhibitory networks. Pup exposure and maternal experience trigger an episode of heightened auditory cortical inhibitory plasticity. For example, GAD67 levels are roughly doubled in the auditory cortex of both wild type and *MeCP2^het^* five days after the birth of the litter. This result points to a reorganization of the cortical GABAergic network triggered by maternal experience. Although this feature of the auditory plasticity is shared between *SurWT* and *SurHet*, *SurHet* also show large increases in expression of PV and PNNs on the fifth day of pup exposure. Notably, initial levels of these inhibitory markers in *NaïveWT* and *NaïveHet*, and levels in surrogates after pups are weaned, are identical. Therefore, potentially crucial features of *MeCP2^+/−^* pathology may only be revealed by the commencement of an episode of heightened sensory and social experience, as occurs with first-time pup exposure. We speculate that this may be a general phenomenon wherein exposure to salient sensory stimuli may define a particularly vulnerable point for *MeCP2^hets^*. Behavioral assessment of other mouse models during similar exposures that challenges network plasticity mechanisms may reveal intensified phenotypes.

### MeCP2 mutation interferes with plasticity by disrupting dynamics of inhibitory networks

Both WT and *MeCP2^het^* female mice exhibit low GAD67 expression as maternally-naïve adults. Expression sharply increases after exposure to a mother and her pups, and returns to baseline levels when the pups are weaned. This is correlated with a surge in the expression of PV and PNNs in *MeCP2^het^* only. This result is consistent with increased PV [36] and PNN expression observed in the developing *MeCP2-null* visual cortex [42]. Several lines of evidence implicate elevated expression of PV and PNN as ‘brakes’ that terminate episodes of plasticity in development and adulthood. In the developing cortex, maturation of GABAergic inhibition mediated by the fast-spiking PV interneuron network is a crucial mechanism for regulating the onset and progression of critical periods [39]. During postnatal development, PV interneurons undergo substantial changes in morphology, connectivity, intrinsic and synaptic properties [53-56], and they form extensive reciprocal chemical and electrical synapses [53, 57, 58]. Learning associated with a range of adult behaviors might rely on similar local circuit mechanism observed in the developing cortex [25, 59].

PNNs inhibit adult experience-dependent plasticity in the visual cortex [24], and in consolidating fear memories in the amygdala [60]. PNN assembly in the *SurHe*t tracks with changes in PV expression after maternal experience, suggesting there is remodeling of the extracellular matrix during natural behavior. This is an interesting observation as the prevailing notion of PNNs during adulthood is as a stable, structural barrier, which needs to be removed with chondroitinase ABC to reactivate plasticity. We propose that impaired experience-dependent cortical plasticity, particularly in PV+ interneurons, contributes to behavioral symptoms in RTT.

Importantly, we demonstrate that manipulating GAD67 expression using *Gad1* heterozygotes is sufficient to restore normal PV and PNN expression patterns and behavior. This result suggests a critical role for *Gad1* in regulating MECP2-mediated experience-driven cellular and circuit operations. MECP2 directly occupies the promoter regions of *Gad1* and *PV* [36, 37], thus potentially configuring chromatin in these promoter and enhancer regions for appropriate activity-and experience-dependent regulation. We speculate that MECP2 regulates specific ensembles of genes and the temporal profile of their expression to control the tempo of plasticity. MECP2 regulates many genes [13, 61]; therefore there are likely other as yet unappreciated targets that could contribute to this control

### Summary

Our data are consistent with an emerging body of literature that suggests that auditory cortical plasticity is triggered in adult female virgin mice by pup exposure. By using pup retrieval behavior as ‘readout’ of the efficacy of this change, we observe that impaired MECP2 expression disrupts both behavior and the underlying auditory cortical plasticity This is consistent with recent data revealing sensory impairments in individuals with RTT, which may contribute to behavioral symptoms [36, 62, 63]. We further speculate that MECP2 deficiency results in experience-dependent “negative” plasticity [64] that may act at other brain regions and time points to contribute to a range of altered behaviors.

## VIDEO LEGEND

**Female *MeCP2^het^* mice failed to retrieve pups on postnatal day 5**. Wild type mother and surrogate WT (*SurWT*) readily retrieve scattered pups, while surrogate *MeCP2^het^* (*SurHet*) fails to gather all pups in a 5-minute session. Video speed for *SurHet* is 8x.

## ACKNOWLEDGEMENTS

The authors wish to thank D. Huang and A. Chandrasekhar for data collection and analysis assistance, and Stephen Hearn at the CSHL microscopy facility for assistance with Volocity software. We would also like to thank A. Zador, B. Li, R. Froemke, J. Tollkuhn, J. Morgan, D. Eckmeier, B. Cazakoff and A. Maffei for helpful comments and discussion.

